# *Toxoplasma gondii*’s basal complex: the other apicomplexan business end is multifunctional

**DOI:** 10.1101/2022.02.23.481603

**Authors:** Marc-Jan Gubbels, David J. P. Ferguson, Sudeshna Saha, Julia D. Romano, Suyog Chavan, Vincent A. Primo, Cynthia Michaud, Isabelle Coppens, Klemens Engelberg

**Affiliations:** Department of Biology, Boston College, Chestnut Hill, MA 02467, USA; Nuffield Department of Clinical Laboratory Science, University of Oxford John Radcliffe Hospital, Oxford, UK; Department of Biological and Medical Sciences, Faculty of Health and Life Science, Oxford Brookes University, Gipsy Lane, Oxford OX3 0BP, UK; Department of Molecular Microbiology and Immunology, Bloomberg School of Public Health, Johns Hopkins University, Baltimore, MD, USA

**Keywords:** *Toxoplasma gondii*, basal complex, MORN1, cell division, endodyogeny, bradyzoite

## Abstract

The Apicomplexa are famously named for their apical complex, a constellation of organelles at their apical end dedicated to invasion of their host cells. In contrast, at the other end of the cell, the basal complex (BC) has been in the limelight since it is much less prominent and specific functions were not immediately obvious. However, in the past decade a staggering array of functions have been associated with the BC, and strides have been made in understanding its structure. Here, these collective insights are supplemented with new data to provide an overview of the understanding of the BC in *Toxoplasma gondii*. The emerging picture is that the BC is a dynamic and multifunctional complex, with a series of (putative) functions. The BC has multiple roles in cell division: it is the site where building blocks are added to the cytoskeleton scaffold; it exerts a two-step stretch and constriction mechanism as contractile ring; and it is key in organelle division. Furthermore, the BC has numerous putative roles in ‘import’, such as the recycling of mother cell remnants, the acquisition of host-derived vesicles, possibly the uptake of lipids derived from the extracellular medium, and the endocytosis of micronemal proteins. The latter process ties the BC to motility, whereas an additional role in motility is conferred by Myosin C. Furthermore, the BC acts on the assembly and/or function of the intravacuolar network, which may directly or indirectly contribute to the establishment of chronic tissue cysts. Here we provide experimental support for molecules acting in several of these processes, and identify several new BC proteins critical to maintaining the cytoplasmic bridge between divided parasites. However, the dispensable nature of many BC components leaves many questions unanswered regarding its function. In conclusion, the BC in *T. gondii* is a dynamic and multifunctional structure at the posterior end of the parasite.

## 1 Introduction

The obligate intracellular Apicomplexa comprise parasites of a wide variety of animal phyla, including humans. All Apicomplexa share a universal body plan designed to invade their next host cell [1; 2]. The structures dedicated to host cell invasion consist of cytoskeleton elements and secretory organelles that are concentrated on the apical end of the cell, which distinct appearance gave the Apicomplexa their name [3]. The other end of the cell, the basal complex (BC), is morphologically much less pronounced, and has received much less attention. The historic appreciation for the BC revolves around its prominent role in cell division: the BC functions as the contractile ring in separating daughter parasites at the conclusion of cell division (**Fig 1**) [4]. The BC is situated at the most basal extremity of the inner membrane complex (IMC), which is the apicomplexan membrane skeleton that orchestrates cell division [5]. The IMC is part of the cortical cytoskeleton that is defined by flattened membrane vesicles (alveoli) decorated on the inside by a set of sub-pellicular microtubules originating at the apical end and a meshwork of intermediate filament-like proteins (alveolins or epiplastins) anchored to the membrane [6; 7]. The cortical cytoskeleton also anchors the myosin motors that power gliding motility and host cell invasion [7; 8]. Indeed one of these motors, MyoB/C, resides at the basal end [9]. Furthermore, the BC maintains a cytoplasmic bridge between parasites and the residual body after division is finalized, facilitating cell-cell communication [10]. Beyond these, putative BC functions are emerging in nutrient uptake [11] and bradyzoite formation and/or maintenance [12; 13] (**Fig 1**). The BC is also the site of intravacuolar network (IVN) assembly, a tubular membrane structure inside the vacuolar compartment required for access to host cell derived vesicles and establishing bradyzoite cysts, though a direct functional involvement of the BC in this process has not been established [14]. Regarding BC dynamics, discrete developmental steps in the BC can be appreciated during the division process [15], whereas in extracellular parasites the very basal end presents as a cup, i.e. the ‘posterior cup’ with a small pore [16]. Recent progress in defining the composition of the BC together with an expanding spectrum of functions has resulted in new insights in the molecular basis of structure-function relationships in the BC, although as discussed here, many questions remain.

**Figure 1.**
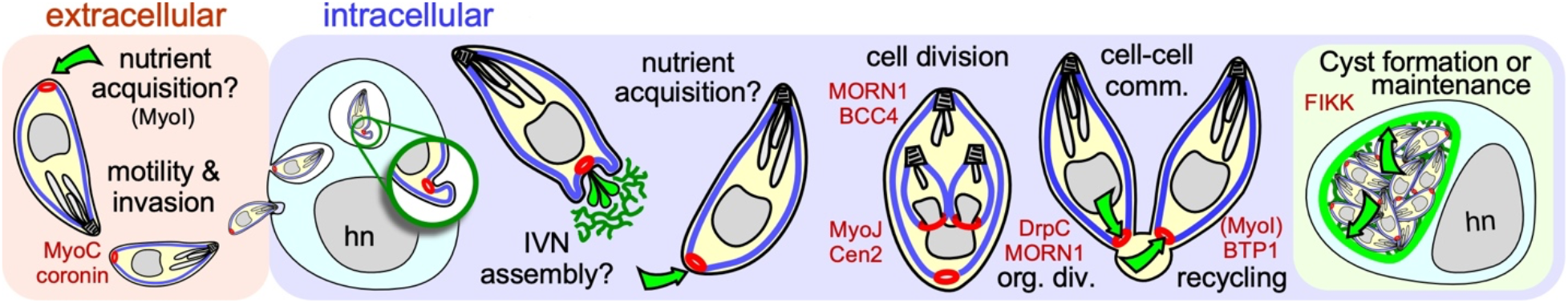
Schematic overview of established and putative roles of the BC in the intermediate host. The BC is represented in red, the IMC in blue, the IVN and cyst walls in green. Green arrows display transport through the BC. Key molecular players are shown in red at the steps where they have key functions. hn marks the host cell nucleus.

## 2 Architecture and Dynamics Throughout the Lytic Cycle

At the ultrastructural level, the BC changes throughout *T. gondii*’s lytic cycle. We differentiate five different arrangements of the BC at the basal end of the cytoskeleton that correlate with its different functions (**Fig 2A**). During the first half of daughter cytoskeleton assembly no electron dense structure is visible at the basal end. However, we identified two proteins that are present at the BC at the very early steps in daughter cytoskeleton formation: the scaffolding protein MORN1, which is first seen as a hazy cloud surrounding the duplicated centrosomes before assembling in a ring at the initiation of each daughter cytoskeleton formation [17; 18; 19] and BCC4, a protein without identifiable functional domains that assembles independently from MORN1 into a ring-like structure [20] (**Fig 2B**). MORN1 and BCC4 are in a complex together and disruption of either protein results in incomplete daughter separation at a very severe fitness cost [20; 21; 22]. These early BC proteins are essential to stabilize the growing basal ends of the daughter buds and recruit additional BC proteins. Several additional proteins associate with the BC while the cytoskeleton scaffolds are growing, including several hypothetical proteins, two phosphatases and microtubule binding protein DIP13 [23] (**Table 1**). Of these, the HAD2a phosphatase is essential for the parasite, and its absence presents a phenotype similar to MORN1 and BCC4 depletion [24], suggesting that it regulates the stability of the BCC4/MORN1 interaction.

**Figure 2.**
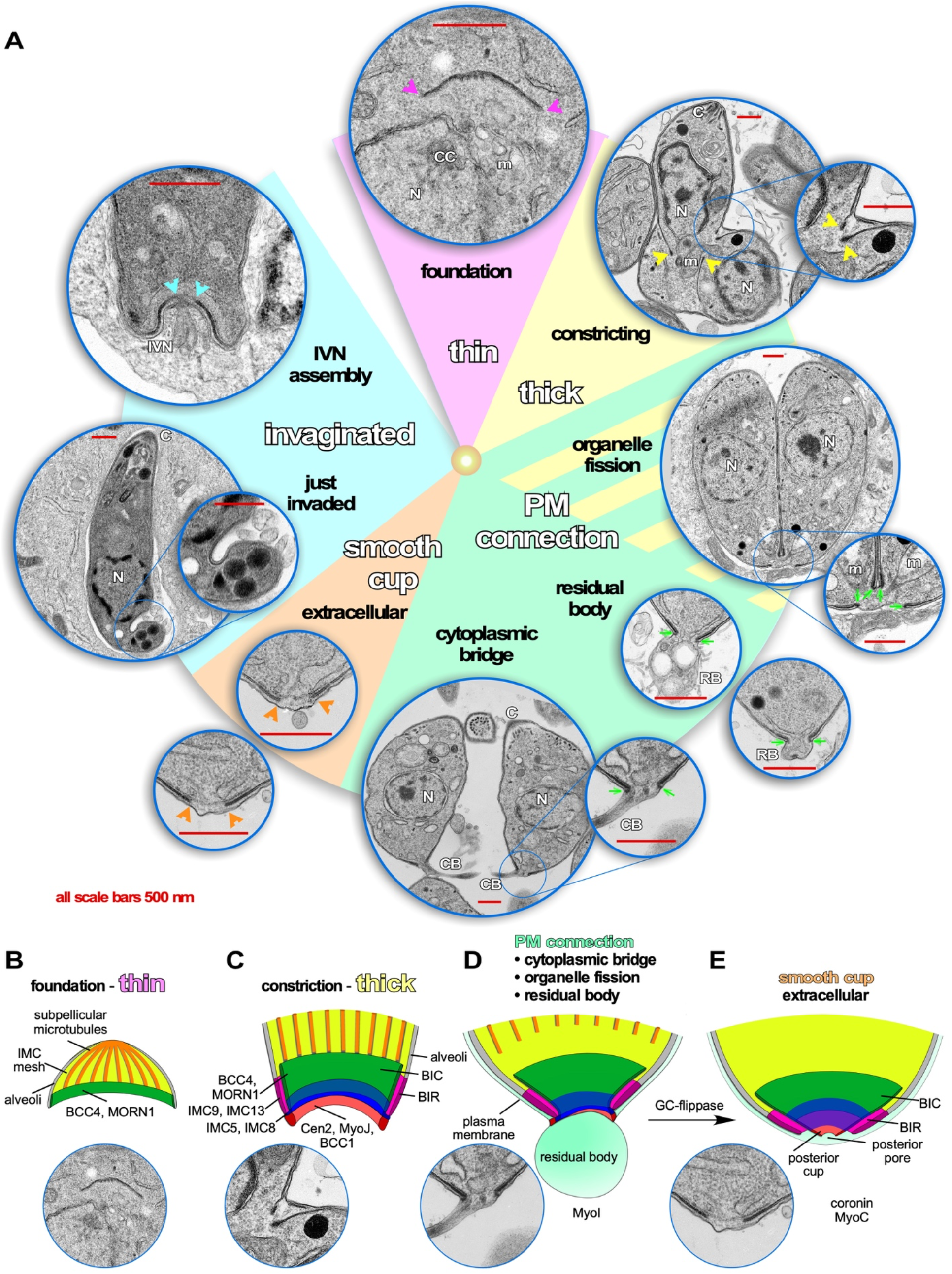
Dynamics of the Basal Complex (BC) throughout the *Toxoplasma* lytic cycle. A. Pie slice colors reflect different manifestations of the BC illustrated by representative transmission electron microscopy (TEM) images. Arrows and arrowheads in the corresponding color point out the BC at critical points. **Purple**: during early daughter bud formation, the basal end of the forming cytoskeleton is thin, although several molecular markers of the BC are present at these early stages such as MORN1 and BCC4, which comprise the foundation for recruitment of additional markers. **Yellow**: when daughter development is halfway, the BC transforms into a double-layer of electron dense material, which coincides with the constriction that tapes the cytoskeleton buds toward the posterior end. **Sea green**: When the plasma membrane fully coats the daughter cytoskeleton, an electron-dense connection from the BC toward the plasma membrane is visible. It is postulated that this connection might be needed to partition the mitochondrion as well as to recycle material from the residual body and/or to maintain the connection with the residual body, as well as to maintain the cytoplasmic bridge connecting tachyzoites within the same vacuole that is responsible for the typical rosette-shaped tachyzoite manifestation. **Orange**: once the cytoplasmic bridge is broken and parasites are extracellular, the BC connection to the plasma membrane disappears and is replaced with a continuous structure on the inside of the IMC. Most likely, this is the “posterior cup” observed in detergent extracted cytoskeletons of tachyzoites [16] (we note that a small hole is present in the extreme end of the detergent extracted posterior cup, which is not observed in the whole parasite TEMs shown here). Endocytosis, likely mediated by coronin that translocates to the BC in extracellular parasites, is required for efficient gliding [33; 35]. **Baby blue**: minutes after completion of host cell invasion by tachyzoites, the basal end of the parasites has a twisted appearance. This is likely the result of tachyzoite rotations within the vacuole to pinch of the PVM [36]. A couple of minutes later, the BC itself invaginates, still maintaining a continuous layer within the IMC, which coincides with the assembly of the IVN from the basal end. However, a direct link between BC appearance, composition or function and IVN assembly has so far not been observed and might merely be coincidental. **B-E.** Schematic representations of the architecture and molecular organization of the four different BC stages recognizable by TEM. **B.** First half of daughter bud assembly. **C.** Second half of daughter bud assembly. **D.** BC in mature but conjoined parasites before abscission of divided cells. **E.** BC in a fully mature, single tachyzoites (motile if extracellular).

**Table 1.**
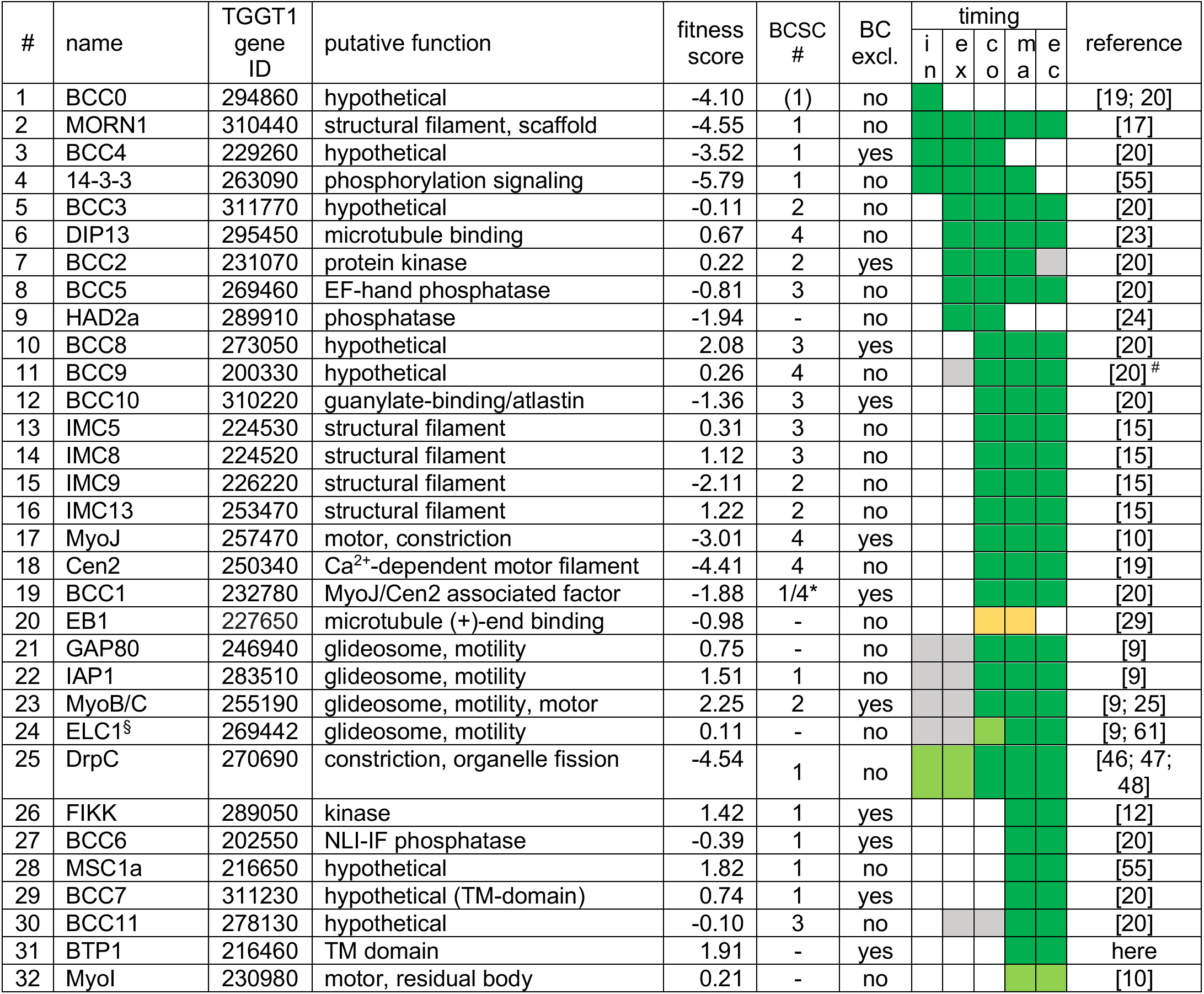
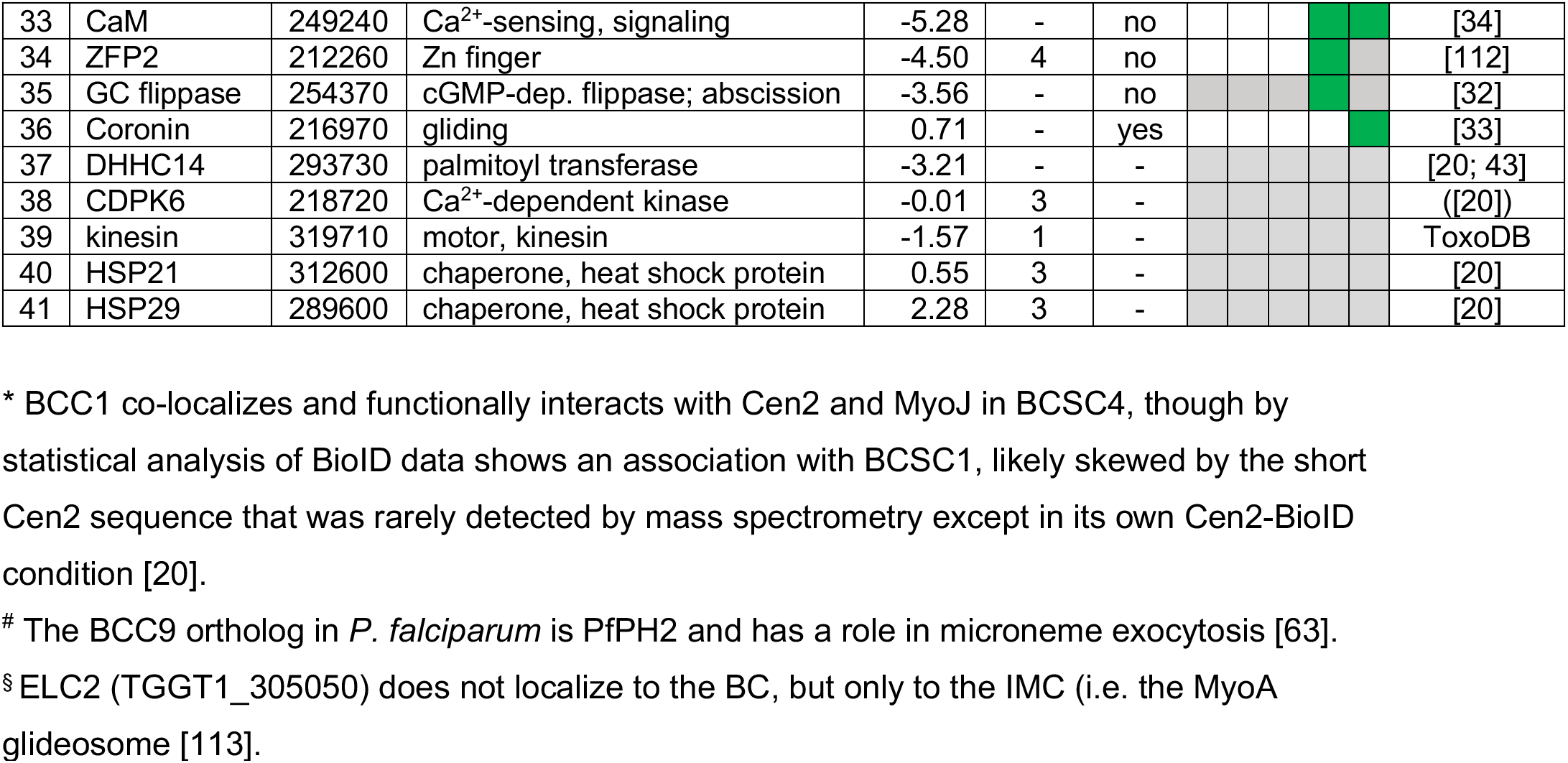
Overview of all known *T. gondii* proteins mapping to the BC. Gene IDs are derived from ToxoDB.org [111]. ‘Putative function’ is based on functional analysis and/or domains contained in the protein as annotated on ToxoDB. ‘Fitness score’ is derived from the genome wide CRISPR/Cas9 KO screen; a score of <-2.5 is a general prediction of gene essentiality [57]. ‘BCSC’ represents the BC sub-complex as defined by structural dissection of the BC by proximity biotinylation [20]. ‘BC excl.’ indicates whether the protein exclusively localized to the BC (yes) or is also seen at other sub-cellular localizations (no). ‘Timing’ of BC association is defined as follows: in, initiation; ex, expansion; co, constriction; ma, maturation; ec, extracellular. Color coded boxes report the following: green: present at BC; white: absent from BC; grey: BC presence not determined; yellow: in the vicinity of the BC, but direct association unlikely; light green (chartreuse): tentative association with the BC.

Halfway through daughter assembly, the BC thickens into an electron dense structure that appears at the basal end of the nascent cytoskeleton scaffolds. At this point the cytoskeletons are being assembled within the cytoplasm of the mother cell (**Fig 2A, C**). Here, the BC constricts, mediated by Myosin J (MyoJ) and Centrin2 (Cen2) [10], that likely work together with a recently discovered component, BCC1 [20]. At the same time, additional cytoskeleton proteins [15] as well as phosphatases and kinases are recruited. MyoC, is deposited at the BC at this point as well, although its key function is much later, in motility of the mature parasite [10; 25; 26]. Over 20 proteins are present at the BC at the time of its constriction, which resolve into four different BC sub complexes (BCSC) based on statistical analysis of proximity biotinylation data (BCSC1-4; **Table 1**; [20]). This mirrors the several substructures discernable by electron microscopy (**Fig 2A, C**). Loss of either MyoJ or Cen2 from the BC results in incomplete BC constriction but has only a modest effect on parasite fitness *in vitro*, suggesting this step is not strictly essential [10]. This observation fits with the modest effect of actin depolymerizers on daughter cell formation [27] and that the deletion of the actin gene does not prevent the completion of cell division [28].

While the BC is constricting, the extension of the sub-pellicular microtubules stops and their (+) ends dissociate from the BC when they are at about 2/3 of the final length of the parasite. At this point the microtubule end binding protein EB1 is briefly visible at the BC, although it is not clear if it directly associates with the complex (**Table 1**; [29]). This event most likely coincides with DIP13 dissociation from the BC [23], and while the mechanism is not well understood, its timing overlaps with completion of karyokinesis, an event in which EB1 has a dynamic role as well [29]. The next BC transition occurs when the plasma membrane is deposited on the IMC: an electron-dense connection from the BC folding around the end of the IMC to the plasma membrane is now formed (**Fig 2A, D**) [15]. Dynamically, this is when the BC pinches to the max. At this point BCC4 releases from the BC [20] (**Table 1**). Functionally, this BC constellation is associated with several different events late in the cell division process: completion of plastid and mitochondrial division and partitioning; depositing of mother remnants into the residual body and their sub-sequent recycling (i.e. re-uptake in the daughters) and finally the maintenance of a cytoplasmic bridge between the daughter parasites upon completion of cell division [10; 28; 30; 31].

Following complete daughter separation, the BC connection from the inside of the IMC to the plasma membrane is lost and the structure transforms into the much more electrolucent posterior cup, which has a small pore at the very basal tip [16] (**Fig 2E**). This complete separation requires the activity of an unique guanylate cyclase (GC) flippase protein present at the BC [32].

Finally, upon egress from the host cell into the extracellular environment, the parasites become motile. MyoB/C contributes to gliding motility and subsequent invasion of the next host cell [9]. In motile parasites, the F-actin stabilizing protein Coronin is recruited to the BC in response to changes in intracellular Ca^2+^ [33], which might be mediated by calmodulin at the BC ([34]; **Fig 3A**). Coronin has also been tangentially associated with microneme secretion, likely due to its putative role in endocytosis occurring at the basal end to offset membrane surplus by microneme secretion at the apical end of the cell [35].

**Figure 3.**
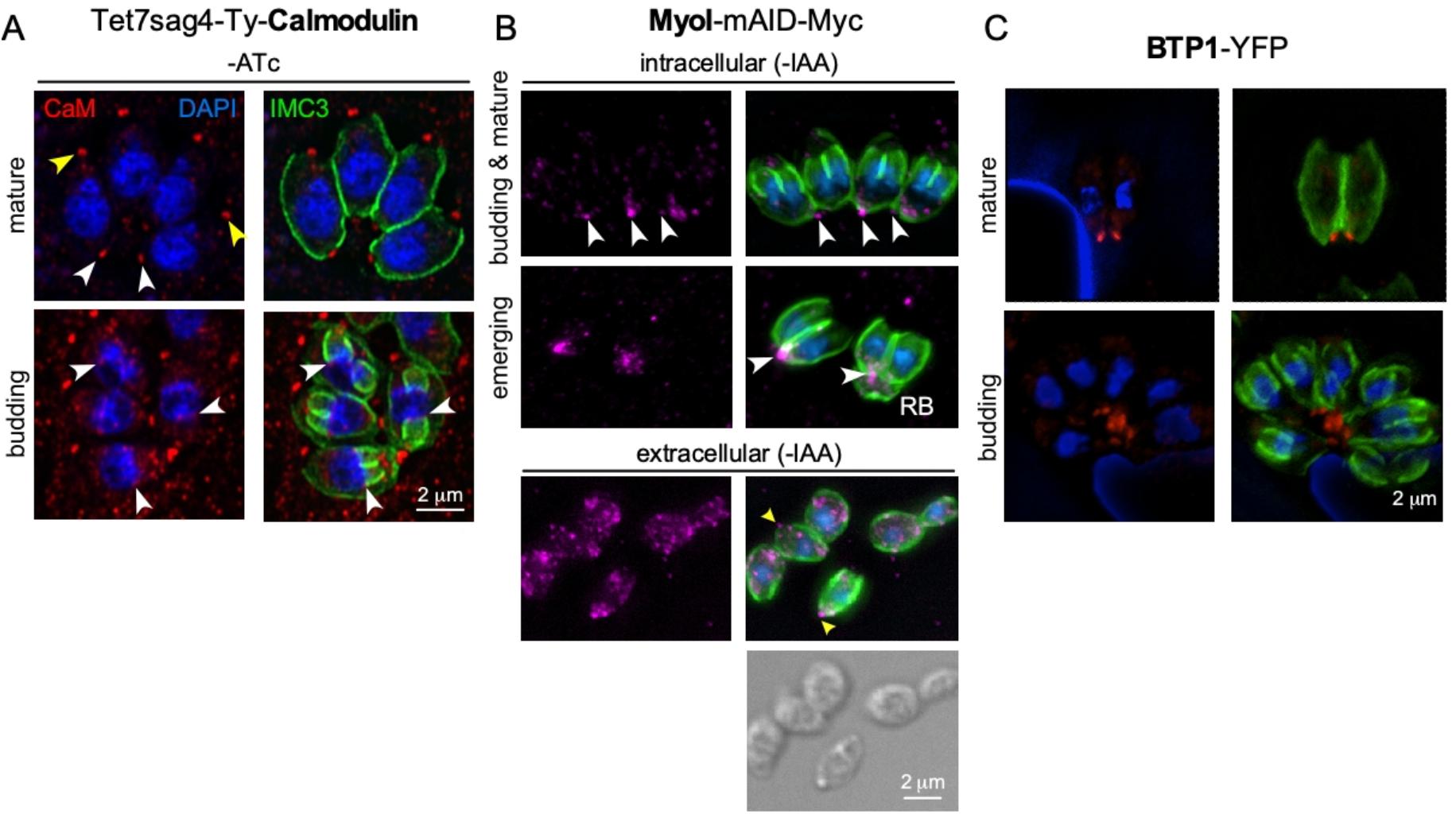
Additional players at the basal complex. **A.** Endogenously tagged Calmodulin localizes to the BC in mature parasites and the daughter buds upon constriction (white arrowheads), next to a more prominent localization at the apical end (yellow arrowheads). IMC3 marks the cortical cytoskeleton of mother and daughter parasites; DAPI marks the DNA. **B.** Endogenously tagged MyoI displays a spotty pattern reminiscent of vesicles. MyoI accumulates in and/or around the BC in mature parasites and in the (forming) residual body (RB), as marked by white arrowheads. In extracellular parasites the signal is more diffuse throughout the cytoplasm but a prominent signal around the BC is observed in many parasites as marked by yellow arrowheads. **C.** Exogenously expressed BTP1 tagged with YFP (false colored red) under its native promoter localizes to the BC in the mature cytoskeleton only. IMC3 marks the cortical cytoskeleton of mother and daughter parasites; DAPI marks the DNA.

Upon conclusion of the host cell invasion process, the parasite twists to seal of the vacuole [36], resulting in a twisted appearance of the basal end (**Fig 2A**, invaginated). It is not clear whether the BC has an active role in this process, or whether the distorted BC appearance might be a physical result of the contortion accompanying twisting. Shortly after completion of invasion the parasite starts to assemble the IVN at the basal end, a process where an active role of BC is yet to be demonstrated (**Fig 2A**) [14]. Thus, current knowledge on the BC during assembly, cell division and cell-cell communication is quite detailed, but our understanding of BC manifestation and function during invasion and the assembly and/or function of the IVN is very limited.

## 3 Cell division

The distinct ultrastructural presentation and key composition changes of the BC during cell division are discussed in the previous section. The prominent role of the BC during cell division is to function as the contractile ring, and to separate the daughters. The mechanism was a puzzle for a long time since depletion of the obvious MyoJ motor complex disabled BC constriction, but only caused a very minor growth phenotype [10], while depletion of BC scaffolding protein MORN1 results in lethal, conjoined daughters unable to separate from each other [21; 22]. This paradox suggested the presence of an additional contractile event preceding the recruitment of the MyoJ/Cen2/BCC1 complex at the midpoint of budding. We recently described a stretchy rubber band model of the BC, where it is the stability of an expandable BC ring with MORN1 and BCC4 as key players, that is the most critical function of the BC during cell division. In absence of either component, the BC is initially assembled, but at the midpoint the complex suddenly breaks into pieces, which results in fraying of the extending sub-pellicular microtubules [20]. This effect is best described as opening an umbrella, and results in daughter buds that have very wide basal diameters, which in the follow prevents their separation (these conjoined daughters have been dubbed the ‘double-headed’ or ‘multi-headed’ phenotype).

Resolving this conundrum was a major breakthrough in understanding the key role of the BC during cell division.

There are several additional phenomena during cell division with crucial roles of the BC. BC formation is first visible as MORN1 and BCC4, which are assembled on a five-fold symmetrical structure. This foundational structure comprises proteins that form the basis for the three distinct major components of the cytoskeleton scaffolds (microtubules, alveolar vesicles, epiplastins/alveolins) [20]. IMC32 and F-BOX ubiquitin ligase (FBXO1) are required for IMC membrane skeleton formation [37; 38], whereas AC9 is critical in conoid and sub-pellicular microtubule formation [39; 40]. In addition, two proteins, BCC0 and BCC3, also exhibit this distinct 5-fold symmetry and were specifically detected in proximity biotinylation experiments using BC components as bait [20]. Further catering to this assessment is that BCC0 has been posited as a potential ortholog of the radial spoke protein 5 (RSP5) in *Chlamydomonas reinhardtii*, which binds to MORN domain proteins, although the statistical support for this homology is moderate at best [41; 42]. Furthermore, BCC3, is deposited in the early BC, from which it transitions into the sutures between the alveolar membrane sheets that make up the IMC. This suggests a direct connection between the five-fold symmetry of the alveolar structure and the BC as well as extending the daughter buds. An additional player in IMC formation is palmitoyl acyltransferase DHHC14 which has been functionally associated with IMC formation [43]. Moreover, DHHC14 is prominently detected in the BC by proximity biotinylation [20], and together these insights support a model wherein the BC serves as the docking site to add new cytoskeletal components by palmitoylating proteins deposited into the alveolar vesicles.

Furthermore, addition of Golgi derived vesicles at the basal end of daughter buds is mediated by Rab11b GTPase [44], which ties the BC into this process. These observations all algin with the apical to basal direction of the daughter assembly and put the BC at the place of daughter bud extension. This model comes with an exception: the apical cap section of the alveolar membrane skeleton appears to bud in the opposite, apical direction away from the five-fold symmetrical foundational structure [20].

Furthermore, induced defects in lipid metabolism, either pharmacologically by ciprofloxacin [45] or genetically by disruption of an apicoplast acetyltransferase (ATS2) [46], result in a cell division stall at the late stage when the mother’s plasma membrane is added and daughter buds start to emerge. Incomplete BC constriction is also observed here, which therefore seems to be tied, directly or indirectly, to the addition of IMC and/or plasma membrane to the emerging daughters. Interestingly, in the ATS2 depleted mutants, DrpC localization to the BC of dividing daughters is defective, which due to its putative membrane constrictive properties was suggested to be causing the lack of constriction ([46]; DrpC docking on the BC was shown to be dependent on the presence of phosphatidic acid). Reciprocally, direct depletion of DrpC does result in two different phenotypes: 1. defective delivery of vesicles to the growing IMC [46; 47]; and 2. incomplete mitochondrion division ([48] further discussed below). Overall, these phenotypes suggest a model wherein fatty acid production in the apicoplast ensures the availability of phosphatidic acid at the BC of budding daughters. This secures the recruitment of DrpC, which in turn is needed to add more membrane to the IMC and/or plasma membrane.

Finally, another pharmacological agent inducing a similar phenotypic block in plasma membrane addition is itraconazole, which at least in fungi inhibits sterol metabolism, and thus most likely affects membrane biogenesis in *T. gondii* [49]. Taken together, vesicular membrane docking at the BC is required to complete cell division.

## 4 Organelle division

Firstly, completion of apicoplast division is dependent upon MORN1 [22]. During apicoplast division, the extended organelle is anchored in the daughter buds by association of the ends to the centrosomes while the undivided organelle is seen extending to the basal ends of the daughter cytoskeleton buds with a sharp bend where the basal complexes meet [17; 50].

Furthermore, the dynamin related protein A (DrpA) is enriched at the pinching point of the apicoplast where the BCs of the daughters come together [51]. However, whether DrpA is specifically recruited to the BC or whether DrpA localizes to the physically most strained plastid region has not been established. In chronological order, apicoplast division completes at the onset of BC constriction, i.e., coinciding with the assembly of the electron dense bulb containing the IMC proteins, MyoJ and Cen2 (**Fig 2C**).

Secondly, the single copy mitochondrion of *T. gondii* is the last organelle entering the daughter buds around the time when the plasma membrane deposition is completed and the residual body is formed (**Fig 1**) [52]. The mitochondria are anchored to the daughter bud IMC through lasso-mitochondria factor 1 (LMF1) [53], whereas their extended membrane structure is directed into the daughter buds in a DrpC dependent fashion [48]. This phenomenon again might be mediated by DrpC-dependent vesicular transport [46; 47]. In conclusion, the BC has been associated with completion of apicoplast segmentation and partitioning of the mitochondrion, but the exact mechanism in both cases is not well resolved.

## 5 Residual body and cytoplasmic bridge

In the late stages of division, when the daughter cells emerge from the mother, remnants from the mother are deposited in the residual body. The residual body sits at the basal end of the daughters and quickly disappears following completing of division (e.g. **Fig 3C**). Thus, the residual body is a recycling bin that is quickly emptied. There are two models on what happens with the residual body and its digested content, which may both be true: 1. The residual body is slowly degraded inside the vacuole and is assimilated into the surrounding IVN tubules; 2. The digested contents are resorbed into the daughters that are still connected to the residual body. The former model could apply if the residual body gets severed from the daughter cells.

However, it is frequently observed that after the residual body is emptied, the connection between daughter parasites, known as the cytoplasmic bridge, is maintained. This ‘cytoplasmic bridge’ facilitates cell-cell communication between parasites and results in synchronized cell division cycles within the vacuole [10; 28; 30]. Upon several sequential division rounds, the cytoplasmic bridge connects multiple parasites in the vacuole and is also responsible to maintain rosette organization of parasites [54]. Since the connection originates at the basal end, the BC is in a key position to function in the residual body and the cytoplasmic bridge. MyoI is an essential player in establishing and/or maintaining the cytoplasmic bridge (**Fig 3C**), with a supportive role assigned to MyoJ [28]. Why the cytoplasmic bridge is maintained is not known as depletion of MyoI does not cause a change in parasite fitness [28]. Although presence of the bridge ensures synchronized division, an obvious need to maintain synchronized cell cycles is not evident. In fact, the bridge is lost in activated macrophages (possibly due to external force acting on the vacuole) and during bradyzoite differentiation, facilitating the asynchronized, slow division in the tissue cysts [10].

We reasoned that besides MyoI, proteins recruited to BC upon completion of cell division might contribute to stabilizing the cytoplasmic bridge (**Table 1**). This list comprises the FIKK kinase [13], CaM (**Fig 3B**; [34]), and MSC1a, a protein of unknown function [55], and two new proteins (BCC6, a phosphatase and BCC7, a hypothetical protein), which were recently added to this list [20]. Furthermore, here we mapped one additional protein: the *T. gondii* ortholog of *P. falciparum* transmembrane protein 1 (PfBTP1), which was linked to acquisition of the plasma membrane by the emerging merozoites [56]. We cloned the genomic sequence of TgBTP1 including the upstream promoter region in-frame with a YFP reporter gene and observed an exclusive localization to the mature BC and to some extend in the residual body (**Fig 3C**).

Collectively, six out of seven proteins association with the BC upon maturation have fitness scores suggesting they are dispensable for in the lytic cycle [57], whereas fitness-conferring CaM is non-exclusively localizing to the BC and likely has other functions that are essential (**Fig S1**) [34]. Indeed, we were able to generate complete knock-outs for all six genes (**Fig S2**), and generated a conditional knock-down line for CaM using the Tet-off system [58]. We first analyzed whether these mutants had a cytoplasmic bridge phenotype by assessing the incidence of parasite organization in rosettes within the vacuole (**Fig 4A.B**). In comparison to wild type parasites where 70% of the vacuoles display a rosette organization, the mutants displayed more diversity; BTP1 and CaM knockout parasites exhibit rosettes in only 15-20% of vacuoles, while for MyoI, BCC7 and FIKK knockouts rosettes were observed in 35-45% of the vacuoles. For BCC6 and MSC1a-depleted parasites, we found no significantly different amount compared to wild type parasite. Thus, these observations strongly suggest that most mature BC proteins support in one way or another the formation and or maintenance of the cytoplasmic bridge.

**Figure 4.**
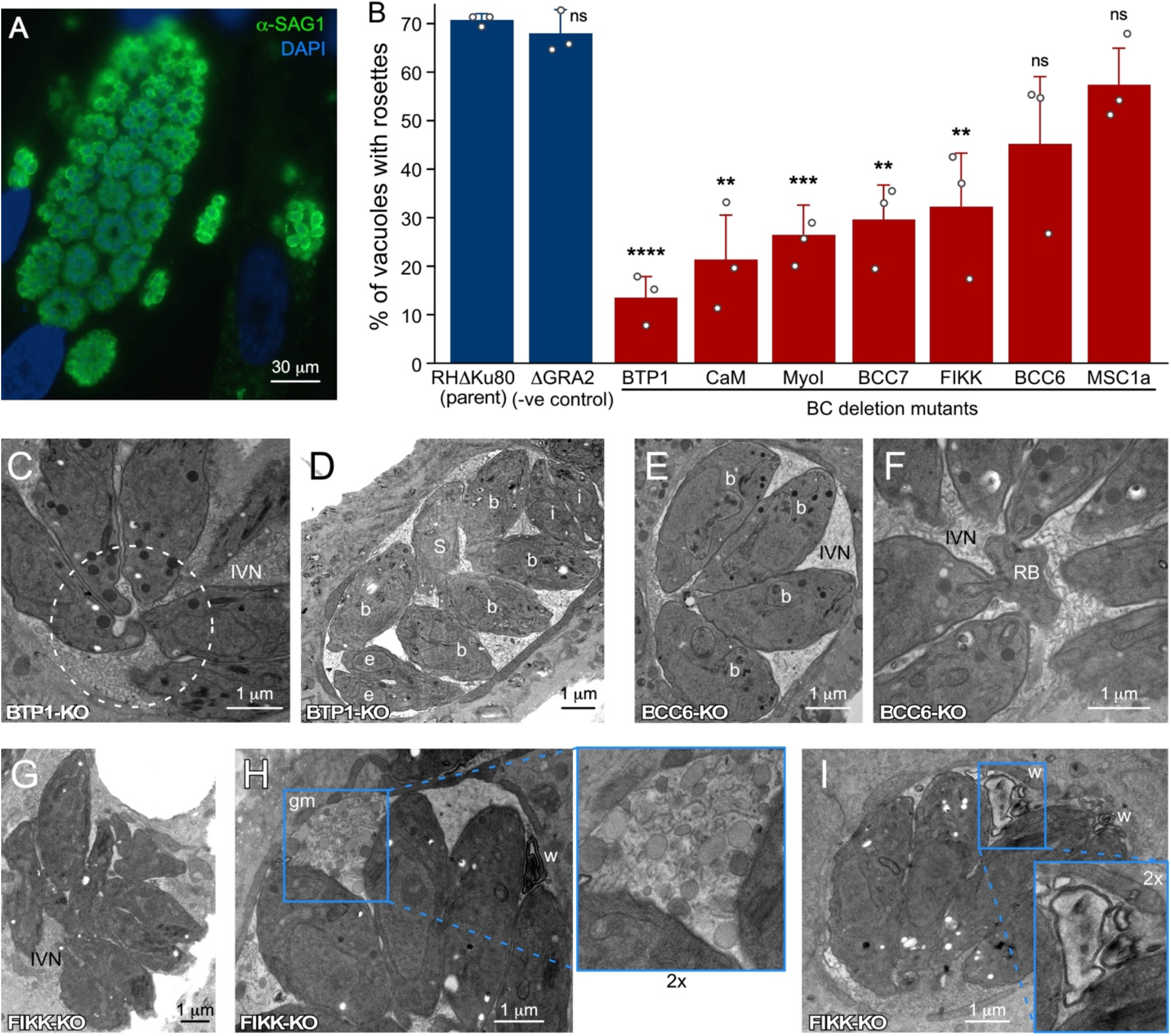
Ultrastructural assessment of the PVM, IVN and intravacuolar organization of parasites wherein genes encoding BC proteins recruited upon completion of division have been knocked out. **A.** IFA example of rosette quantification assay. Shown is the parent line control (RH*Δ*Ku80). **B.** Incidence of parasites organized in rosettes. Only vacuoles with >16 parasites per vacuole were counted, and at least 100 vacuoles per biological repetition were counted. Bars represent the average of three biological repetitions, error bars report standard deviation. Student’s *t*-test relative to RH*Δ*Ku80: * < 0.05; ** < 0.01; *** < 0.001; **** < 0.0001; ns: not significant. **C, D.** BTP1-KO parasites are not connected by a cytoplasmic bridge or to the residual body (C; dotted circle) and divide asynchronously (D; b=budding; e=emerging; S=S-phase; i=interphase/mature), but display normal IVN formation. **E, F.** BCC6-KO parasites show no defects at all and are shown as reference for the wild type manifestation of the IVN, synchrony in cell division, and connections of the basal ends to the residual body. **G-I.** FIKK-KO parasites are not well-organized in the PV (G), display abundant ‘gray matter (gm)’ inside the vacuole of unknow origin or composition (I), and frequently display electron- dense membrane whirls (w) inside the vacuole (H, I). Parasite development stages are marked as follows: b, internal budding; e, emerging daughters; s, S-phase; i, interphase. IVN, intravacuolar network; RB, residual body; gm, gray matter; w, membrane whirls inside the PV.

Subsequently, we dissected representative KO mutant parasite strains at the ultrastructural level (**Fig 4C-I**). This confirmed that BTP1-KO parasites lost the cytoplasmic bridge (**Fig 4C**), and as a result displayed vacuoles harboring non-synchronously dividing parasites (**Fig 4D**).

BCC6-KO on the other hand readily showed synchronously dividing parasites (**Fig 4E**) and connections to the residual body (**Fig 4F**), consistent with the high incidence of rosettes (**Fig 4B**). FIKK-KO organization reflected the rosette assay (**Fig 4G**), but the vacuoles displayed abnormalities in IVN membrane organization such as undefined globules gray matter (**Fig 4H**) and membrane whirls (**Fig 4I**). This suggests that the proteins associating with the mature BC could also impact IVN formation and/or function.

In conclusion, the most prevalent phenotype seen in parasites depleted of proteins associating with the mature BC is the loss of the cytoplasmic bridge and associated secondary phenotypes, such as inability to form rosettes or divide in synchrony. Since we were unable to determine any fitness consequences for the mutant parasites under our experimental conditions, the functional relevance of these numerous BC proteins is currently unclear. It is of note that depletion of LMF1, which anchors the mitochondria in the IMC, also interferes with rosette formation, which might suggest that the cytoplasmic bridge is maintained to accommodate the mitochondrion [59].

## 6 Motility

MyoC resides at the site of the BC and is already seen at the BC halfway during the division process (**Table 1**; MyoB is splice variant of the same gene [25]). However, MyoC does not function in cell division but is involved in gliding motility and host cell invasion [9]. It can replace the function of another class XIV myosin, MyoA, which however alters the tachyzoite’s invasion process and efficiency [60]. MyoC is restricted to the posterior end of the cell by IMC-associated protein 1 (IAP1) and is in complex with glideosome associated protein 80 (GAP80), which forms a bridge between the BC and plasma membrane [9], and Ca^2+^-binding essential light chain 1 (ELC1), but not ELC2 [9; 61]. Thus, the MyoC complex does not reside on the cytoplasmic side of the IMC, but on the outside facing the plasma membrane. The reason for a specialized MyoC at the BC is not known as its depletion has no impact on viability and MyoC orthologs are not widely conserved within the Apicomplexa [9]. It is possible that MyoC is essential in other developmental stages or specialized motility modes, which have not yet been probed.

Another distinct role for the BC in motility is the presence of actin filament binding coronin. Coronin translocation from the cytoplasm to the BC only occurs in extracellular parasites, is Ca^2+^-dependent, and is correlated, likely in a co-dependent fashion, with microneme protein discharge [33]. Recently, motility was associated with the need of endocytic activity in extracellular parasites, which most logically would occur at the posterior end of the parasites, possibly the BC [35]. Thus, coronin-mediated endocytic activity at the BC could be required for efficient motility. Interestingly, *Plasmodium* coronin has also been associated with parasite endocytosis [32] and supports this assignment, although this assignment was made in a different context (hemoglobin uptake rather than motility). Furthermore, MyoI has been shown to be critical for extracellular survival [62] and might be associated with this process as well.

Indeed MyoI is frequently observed at or around the BC (**Fig 3B**) [10]. An additional line of evidence is found in the TgBCC9 ortholog in *P. falciparum*, named PfPH2, which was associated with a role in microneme exocytosis [63; 64]. As evident from the data assembled here, endocytosis of extracellular parasites might play out at the basal end rather than at the apical end, and could be mechanistically related to the microneme secretion defect observed upon coronin depletion in *T. gondii* [33]. Additional work is needed to conclusively tie these processes together at the BC of the parasite.

## 7 PVM and IVN formation

In the last steps of the host cell invasion process the parasite has to seal off the host cell’s plasma membrane, as well as seal the forming parasitophorous vacuolar membrane (PVM). Recent work has shown that a twisting motion by the parasite inside the vacuole mechanically induces host cell membrane fission and PVM sealing to complete the invasion within a protective vacuole [36]. Since this sealing occurs at the posterior end of the parasites, which is the last part of the parasite to enter the cell and PVM, the posterior end of the parasites often appears twisted in newly invaded parasites (**Fig 1, 2A**). In addition, contorted parasites are sometimes also seen among parasites within larger vacuoles (e.g. the far left parasite in **Fig 4C**, or as reported in [65]). Although the BC appears to be at the center of the action here, there are currently no experimental data to support a direct and/or active role.

Another event that unfolds at the posterior end of the parasite, specifically at the site of the BC, 10-15 min post completion of invasion is the formation of the intravacuolar tubulovesicular network (IVN; a.k.a tubulo-vesicular network or TVN) (**Fig 1, 2A**). The IVN is composed of a membranous interface derived from multi-lamellar vesicles secreted by the parasite [14; 66] and lipids scavenged from the host [67]. The IVN is stabilized by various dense granule proteins (GRA2 and GRA6) secreted by the parasite [68; 69; 70]. Deletion of GRA2 and/or GRA6 results in loss of the IVN [69] and subsequent poor acute virulence in mice [71; 72] (likely due to impairment of the CD8 response [73]), and markedly reduced chronic infection [74]. However, “how” the IVN is established and maintained is far from resolved. Notably, the debate regarding the site of dense granule secretion in *Toxoplasma* is not settled either. For example, different dense granule populations with different contents exist, and not all may be secreted at the same time, site, or by the same mechanism [75]. Although there is evidence that the dense granules are secreted somewhere at the apical end of the parasite, possibly the through the apical annuli [66; 76], their secretion has also been observed from a specialized invagination at the posterior end of the parasite where the BC resides [14]. Although it was initially reported that GRA2 is released from the posterior end of the parasite as several multilamellar vesicles emanating from the invaginated BC area, another study reports that GRA2 is secreted apically and then recruited to the posterior end [77]. In either scenario GRA2 multilamellar vesicles are deposited in the vacuole at the site of the invaginated BC, which are subsequently assembled into tubulated structures making up the mature IVN [14] (**Fig 1, 2A**). We note that a block of all dense granule secretion would abrogate the secretion of GRA17 and GRA23, which form a pore in the vacuolar membrane and their deletion results in distinctly, osmotically swollen vacuoles [78]. For the BC mutants tested we have not observed such swollen vacuoles and as such our data so far do not support a role of the BC in dense granule secretion.

During IVN formation the BC has an ‘inside-out’ appearance and a direct role for the BC is quite likely, at least for structural support, but direct evidence is absent. We reasoned that BC proteins recruited to the BC following completion of division could potentially function in this process. As mentioned, neither BTP1 nor BCC6 mutants displayed any defects (**Fig 4C-F**).

However, FIKK-KO parasites show IVN abnormalities (**Fig 4G-I**): 1. abundant ‘gray matter’ inside the vacuole of unknown origin or composition, which is sometimes also seen in wild type parasites; 2. presence of electron-dense membrane whirls inside the vacuole, also of unknown origin. In summary, none of the three tested mutants affected IVN formation, but loss of FIKK leads to abnormal structures, which at the very least does associate the BC with processes within the vacuole.

## 8 Nutrient acquisition

Intracellularly residing parasites can acquire nutrients from the host cell in several ways [11; 78; 79; 80; 81; 82]. Germane to a role of the BC in this process is the observation that host derived vesicles accumulate in the parasitophorous vacuole (PV). Specifically, host GFP-Rab11A decorated vesicles accumulate in the center of the PV in the middle of the rosette at the basal ends of the replicating parasites [11], strongly suggesting the participation of the BC in host vesicle remodeling and/or lipid uptake [11] (**Fig 5A**). We utilized our set of parasite BC gene KO strains to determine any changes of host GFP-Rab11A accumulation patterns. As a positive control we generated a GRA2-KO mutant, which in a different genetic background strain was previously shown to abrogate IVN formation and to be critical to this process [11]. GRA2 depletion indeed resulted in an increased distance between GFP-Rab11A vesicles in the PVM and the center of the vacuole (**Fig 5B**) and a decrease of the total number of vesicles observed per vacuole (**Fig 5C**). However, none of the BC mutants show significant changes in foci distance from the vacuole center, and only the FIKK-depleted mutant showed an increase in number of vesicles per parasite (**Fig 5B, C**). Thus, FIKK depletion leads to an accumulation of vesicles in the PV, which is most likely the result of the aberrant membrane whirls and gray matter forming in the vacuole upon FIKK depletion.

**Figure 5.**
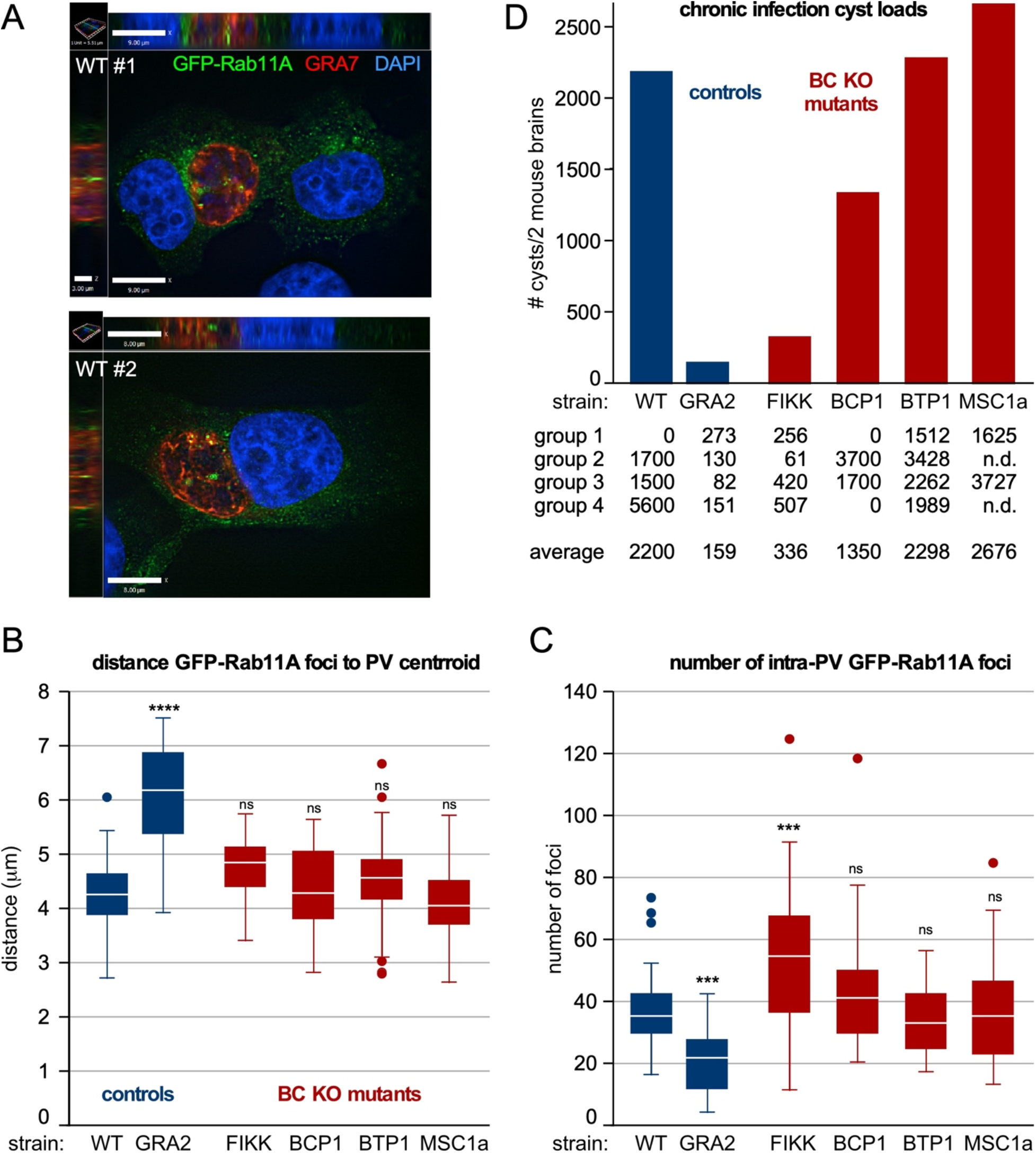
The role of BC proteins recruited upon completion of division in IVN function and establishment of chronic infection. **A-C.** RH strain parasites with BC protein gene knock-outs and controls were inoculated in host cells expressing Rab11A-GFP, which accumulate in the center of the vacuole where the basal ends are clustered [11]. GRA2-KO parasites do not form an IVN and were used as negative control. Panel **A** shows two examples of XYZ projections on which measurements were performed. Shown are wild type parasites infected samples stained additionally with GRA7 to highlight the parasitophorous vacuole membrane and DAPI. **B.** Measurements of distance of the Rab11A-GFP foci inside the vacuole to the centroid using Volocity software. **C.** The total number of foci per vacuole was counted. For B and C, at least 26 vacuoles were counted; box plots depict average ±SD with whiskers representing the upper and lower values excluding outliers; outliers are marked as open circles, and the white line inside the box is the median. ANOVA relative to RH*Δ*Ku80: * < 0.05; ** < 0.01; *** < 0.001; *** < 0.0001; ns: not significant. **D.** Prugniaud strain BC protein knock-out and control parasites were i.p. injected in female C57/BL6 mice (group 1 and 3: 500 tachyzoites; group 2 and 4: 2500 tachyzoites). Brains were harvested 3-4 weeks post infection and cysts purified from two pooled mice brains and enumerated following DBA staining of the cyst wall.

We recently discovered a role for MyoI in extracellular parasites, as increased MyoI expression was associated with prolonged extracellular survival capacity [62]. Moreover, extracellular survival capacity was functionally mapped to the availability of fatty acids. As such, a plausible model is that lipid uptake by extracellular parasites is mediated by MyoI at the BC (**Fig 3B**). Furthermore, uptake of extracellular material would be consistent with endocytosis during gliding, but whether the mechanism is the same remains to be directly tested.

## 9 Tissue cyst formation

All insights discussed so far pertain to the tachyzoite life stage of *Toxoplasma*. However, insights have been emerging for a role of the BC in chronic infection of mice, which is established by differentiation of tachyzoites into bradyzoites. Bradyzoites are not very proliferative or metabolically active and reside within tissue cysts contained by a proteoglycan cyst wall [83; 84]. A putative role for the BC in bradyzoites was revealed by the identification of FIKK mutant that resulted in sharply reduced chronic infection in a mouse model [13].

Furthermore, cyst formation is dependent on the IVN as disruption of key GRA proteins (e.g. GRA2) disrupts IVN assembly and severely impacts the ability to establish chronic infection [74]. It is important to note that the bradyzoite IVN is different from the discrete tubules seen in tachyzoites: the vacuolar space in bradyzoites is a matrix of differently sized vesicles and tubules organized in a network that connects the bradyzoites to each other and to the cyst wall. Moreover, this matrix is involved in the transport of materials from the bradyzoites to the cyst wall [84; 85; 86]. The BC could therefore have a putative role in cyst maintenance and/or function of the matrix. We set out to test whether proteins recruited to the mature BC affect the ability to establish and/or retain brain cysts in mice. Since the already generated KO mutants were in the Type 1 genotype RH strain that is hypervirulent in mice and does not permit the formation of cysts, we generated direct KOs of FIKK, BCC6, BTP1, and MSC1a genes, while GRA2 was used as a positive control in the genotype II Prugniaud (Pru) parasite strain. As expected, the cyst load was severely reduced in both GRA2-KO and FIKK-KO infected mice (**Fig 5D**). However, none of the other mutants displayed a strong reduction in cyst loads. Thus, we conclude that amongst the proteins recruited to the mature BC, only FIKK impacts chronic infection, and hence, the other tested BC proteins are not involved in cyst formation in the bradyzoite stage.

## 10 Outlook

In summary, the BC is a multifunctional structure with several validated and a series of putative functions. Regarding cell division, the BC is the site where building blocks are added to the cytoskeleton scaffold, the BC has a two-step mechanism (stretch and constriction) as contractile ring, and the BC is key in organelle division. Furthermore, the BC has roles in ‘import’, i.e. the acquisition of host-derived vesicles, possibly the acquisition of extracellular lipids and the endocytosis of microneme proteins to facilitate motility. In parallel, the BC is also tied to motility through the presence of MyoC. Furthermore, the BC acts on IVN assembly in a poorly understood fashion, that may directly or indirectly contribute to the establishment and/or maintenance of chronic tissue cysts.

The most remarkable aspect regarding the BC as contractile ring in cell division is that a motor protein is strictly obsolete. Notably, other apicomplexans like *Plasmodium falciparum* seem to have done away with it altogether [87; 88]. Indeed, many protozoa complete cell division without an actinomyosin ring [89; 90]. Since the apicomplexan cell division process shares many features with the assembly of cilia or flagella [42], we hypothesize that the actinomyosin ring together with Cen2 has evolutionary shared ancestry with “transition zone” at the base of motile cilia, sensory cilia and the connecting cilium in photoreceptor cells [91].

Particularly in photoreceptor cells, centrin/G-protein complexes organize signaling in retinal photoreceptor cells, next to a function as a barrier and facilitate transport of vesicles [91]. In Apicomplexa this function may still be intact (e.g. in recycling material from the residual body and/or mitochondrion partitioning), but appears to have been repurposed for additional roles in cell division.

Studies on the BC in *P. falciparum* have identified as set of proteins that is largely not conserved in *T. gondii* [88]. A notable player is PfCINCH, a dynamin-like protein with an essential role in *P. falciparum* BC constriction. In *T. gondii* DrpC is present in the BC but is involved in transport rather than BC constriction [47; 48]. A putative ortholog of PfCINCH is BCC10 as it displays homology with atlastin: in vertebrates atlastin is a dynamin-like GTPase required for homotypic fusion of endoplasmic membrane tubules [92]. However, we did not identify an orthologous protein of BCC10 in *P. falciparum* [20], whereas the mild fitness score indicates BCC10 is not essential, suggesting it is not a functional PfCINCH ortholog. These examples are part of a bigger picture: only very few of the BC components mapped in *T. gondii* and *P. falciparum* are conserved, and if they are conserved, their functions diverged (e.g. MORN1 is widely conserved across division modes [93], but unlike TgMORN1, PfMORN1 is not essential [94]). Another example is the single FIKK gene in *T. gondii*, which contrasts sharply with the expanded FIKK family in *P. falciparum* with functions in non-cell cycle related processes outside the parasite boundaries [95; 96]. This is puzzling, and possible scenarios are that these proteins are very fast evolving. This is supported by the many coiled-coil regions as the only discernable feature in many the BCC proteins. At the same time, these regions might be reflective of the fundamentally different budding strategy in *T. gondii* vs. *P. falciparum*: internal budding in the cytoplasm of the mother vs. external budding outward at the plasma membrane of the mother cell, respectively [5]. Further work is needed to answer these questions.

Besides the architectural and functional divergence of the BC within the Apicomplexa, there are several additional open questions regarding specific *T. gondii* BC proteins and their functions. The first one is the puzzling set of BC proteins only acquired upon completion of cell division. We show that several function in maintenance of the cytoplasmic bridge, but this bridge is not essential, at least not in tachyzoites. Although we hypothesized these might be involved in import, genetic dissection of the components so far did not identify strong support for this hypothesis. The pursuit of the alternative hypothesis that these proteins function in IVN assembly did not find much support, besides a minor role for FIKK. However, we did confirm the previous observation that FIKK is key to establish chronic infections in mice [13], but none of the other BC proteins in this group contributed to this process.

The disruption of FIKK does not abolish IVN formation, yet sharply reduces the cyst loads. Notably, a similar phenotype was reported for GRA12 depleted parasites, which display an apparently intact IVN but exhibit a delay in the accumulation of the CST1 major cyst wall protein at the cyst periphery [97]. Involvement of the IVN in stage differentiation is unlikely to be related to nutrient acquisition since differentiation in response to nutrient stress is well documented [98; 99]. Since the IVN as seen in tachyzoites presents very different in bradyzoites, as a conglomerate of vesicles and tubules, its function might be different [84; 85; 86]. A putative role for the BC in cyst maintenance and/or function of this network might be dependent on the BC, but beyond FIKK, we found no further experimental support for this scenario. As such, the critical function of most BC proteins associated with mature parasites remains largely unknown.

Another dimension of the BC that is till shrouded in many questions is how its elaborate architecture relates to the variety of functions ascribed to the BC. As shown in **Fig 2** and **Table 1**, BC structure and composition changes significantly throughout tachyzoite development. An established structure-function is that the final constriction requires MyoJ/Cen2/BCC1, which resides at the very basal end of the BC (BCSC-4). Since onset of constriction coincides with many additional proteins being recruited across the various BC subcomplexes, and that these genes are largely non-essential, it is tempting to assign a function buttressing the unessential process of final constriction. On the other hand, these proteins may also be required for recruiting the BC proteins after completion of cell division, which are non-essential in tachyzoites. However, the absence of essential proteins and BC functions complicates determining the nature of structure function relationships. Assuming these structures and function have been retained under selective pressure, it seems that we have not yet identified the relevant pressures. This also raises the question on how these transitions in composition and function are controlled. We mapped a number of kinases and phosphatases (**Table 1**).

Their genome wide CRISPR/Cas9 fitness scores suggest non-essential roles and only HAD2a has a critical function in getting the BC beyond the midpoint likely acting on BCC4-MORN1 stability [24]. It is conceivable that the non-essential kinases and phosphatases are redundant and function in parallel pathways, a scenario that deserves serious consideration as the BC has many critical functions to the parasite whose integrity might be worth the cost of dedicating multiple layers of control. With BC composition and architecture largely mapped in both *T. gondii* and *P. falciparum* the toolboxes are in place to further probe in these open questions to reveal the remaining secrets of the BC.

## 11 Material and Methods

### Parasites and Host Cells

*T. gondii* tachyzoites were maintained and studied in human foreskin fibroblasts (HFF) or studied in VERO cells, as previously described [100]. Host cells were maintained in DMEM media containing 10% serum. *Toxoplasma* strains RHΔKu80ΔHXGPRT [101], RH-TaTiΔKu80 [102], RHΔKu80ΔHXGPRT-Tir1 [103] and PrugniaudΔKu80 (PruΔKu80) [74] were used as the basis for all mutants in this study. The RHΔMyoI line was generated the same way as the GT1ΔMyoI line described before [62]. Stable transgenics were obtained by selection under 1 µM pyrimethamine, 25 µg/ml mycophenolic acid (MPA) combined with 50 µg/ml xanthine or 20 µM chloramphenicol and cloned by limiting dilution.

### Plasmid cloning and transgenic parasite generation

All oligomers used in this study are listed in **Table S1**. All transgenic lines were cloned by limiting dilution and the genotype validated by diagnostic PCRs (**Fig S2**).

The annotated BTP1 encoding sequence was PCR amplified from RH genomic DNA, including 1.5 kb upstream of the start codon annotated on ToxoDB serving as its own promoter, and was cloned into ptub-YFP-YFP(MCS)/sagCAT [15] using *Pme*I and *Avr*II restriction enzymes to replace ptub-YFP. The resulting plasmid was transiently transfected into wild type RH parasites.

To replace the ORF of a gene of interest we first designed CRISPR-Cas9 plasmids that specifically target the 5’ region (and 3’ region in case of FIKK and BCC6) of the respective genes. Oligomers encoding single-guide RNAs were hybridized and ligated into the *Bsa*I- digested pU6-Universal plasmid [57]. To facilitate homologous directed repair we PCR-amplified a resistance cassette that drives either DHFR-TSm2m3 (for KOs done in RHΔKu80) or HXGPRT (for KOs done in PRUΔKu80) under the dhfr promoter sequence. Specific integration was facilitated by inclusion of 35 bp flanks on the 5’ and 3’ end of the PCR amplicon, which are homologous to the side of Cas9 double strand break. For transfections; 40 ug of a single Cas9 plasmid (or 20 ug of each in case two Cas9 plasmids were transfected) was mixed with the PCR amplicon, transfected and parasites selected with the appropriate drug for proper integration of the resistance marker and deletion of the target gene.

Endogenous tagging of MyoI with a mAID-Myc tag was achieved by PCR amplification of a 833 bp genomic DNA fragment 3’ of the MyoI translational start, and cloned by Gibson assembly into into the AAP4-3xMyc-DHFR plasmid [104] in which the 3xMyc tag was replaced with the mAID-3xMyc coding sequence digested with *Pme*I and *Avr*II, Before transfection the plasmid was linearized with *Avr*II, which cuts within the cloned 3’-region to facilitate homologous recombination in RHΔKu80ΔHXGPRT-Tir1 parasites.

To generate ATc-regulatable CaM (TGGT1_249240) expressing parasites, a 1.3 kb genomic DNA fragment downstream of the ATG codon was amplified using primer pairs CaM- BglII-F/CaM-NotI-R, and cloned by *Bgl*II/*Not*I into an N-terminal cMyc/Ty-epitope tagged plasmid derived from the single homologous recombination plasmid DHFR-tetO7Sag4-Nt-GOI (kindly provided by Dr. Wassim Daher, Université de Montpellier I et II; [105]). The plasmid was linearized with *Nar*I before transfection.

### Western blots

Western blot was performed with lysates from parasites treated ±ATc for indicated periods of time. A 12% NuPAGE Bis-Tris (for CaM) (Invitrogen) was loaded with samples prepared by lysis with 1% SDS in 150 mM NaCl and 50 mM Tris-HCl, pH 8.0, of equal numbers of parasites for each experimental condition. Following SDS-PAGE, proteins were transferred to a PVDF membrane (Bio-Rad) and blocked using 5% milk. Blots were probed with mouse α-tubulin MAb 12G10 (1:2000) and mouse α-Ty (1:500; kindly provided by Dr. Chris de Graffenried, Brown University) followed by probing with horse radish peroxidase (HRP)-conjugated α-mouse (1:10000) (Santa-Cruz Biotech) and detection of signal after chemiluminescent HRP substrate (Millipore) treatment.

### Plaque assay

T12.5 culture flasks or 6-well culture plate confluent with HFF cells were inoculated with 100 parasites of choice and grown for 7, 14 or 21 days. Following incubation, the monolayer was fixed with 100% ethanol for 10 minutes and stained with crystal violet [100].

### Indirect Immunofluorescence Assays

For intracellular localization, parasites were inoculated into 6-well plate having coverslips confluent with HFF cells. Following overnight incubation, parasites were fixed with 100% methanol. For extracellular localization, freshly lysed parasites were filtered, pelleted and resuspended in PBS. Thereafter, parasites were added to poly-L-lysine coated cover-slips and allowed to incubate for 30 min at 4°C prior to fixation with 100% methanol. 1% BSA fraction V in PBS was used as blocking agent.

The following primary antisera were used: α-Myc MAb 9E10 (1:50) (Santa-Cruz Biotech), mouse α-Ty (1:500; kindly provided by Chri s de Graffenried, Brown University), rabbit α- IMC3(1-120) (1:2,000 [15]), mouse α-GFP (Abgent; 1:500). Guinea pig α-AAP4 (1:200 [104]).

Alexa 488 (A488) or A594 conjugated goat α-mouse or α-rabbit were used as secondary antibodies (1:500) (Invitrogen). DNA was stained with 4’,6-diamidino-2-phenylindole (DAPI). A Zeiss Axiovert 200 M wide-field fluorescence microscope equipped with a α-Plan-Fluar 100x/1.3 NA and 100x/1.45 NA oil objectives and a Hamamatsu C4742-95 CCD camera was used to collect images, which were deconvolved and adjusted for phase contrast using Volocity software (Perkin Elmer).

### Transmission Electron Microscopy

For basal complex development stage analysis, HFF infected cells were fixed in 4% glutaraldehyde in 0.1 M phosphate buffer pH 7.4 and processed for routine electron microscopy [106]. Briefly, cells were post-fixed in osmium tetroxide, and treated with uranyl acetate prior to dehydration in ethanol, treatment with propylene oxide, and embedding in Spurr’s epoxy resin. Thin sections were stained with uranyl acetate and lead citrate prior to examination with a JEOL 1200EX electron microscope.

Basal complex mutant infected cells were prepared for ultrastructural observations by fixation in 2.5% glutaraldehyde in 0.1 mM sodium cacodylate (EMS) and processed as described [107]. Ultrathin sections were stained before examination with a Philips CM120 EM (Eindhoven, the Netherlands) under 80 kV.

### Rab11A-GFP vesicle assay

VERO cells stably expressing GFP-Rab11A were infected as described before [11]. In brief, infected cells were fixed in PBS with 0.02% glutaraldehyde (EM grade; EMS) and 4% paraformaldehyde and permeabilized with 0.3% Triton X-100 in PBS for 5 min. Cells were blocked with 3% BSA in PBS followed by incubation in *α*-TgGRA7 antiserum [81], washed with PBS, incubated with Alexa conjugated secondary antibody, washed with PBS and then incubated in 1 μg/ml DAPI in PBS followed by PBS washes. Coverslips were mounted on slides with ProLong antifade mounting solution (Alexa secondary antibodies). Optical z-sections of infected cells with PVs containing 16 parasites to normalize the data for PV size were collected using a Zeiss AxioImager M2 fluorescence microscope equipped with an oil-immersion Zeiss plan Apo 100x/NA1.4 objective and a Hamamatsu ORCA-R2 camera. Optical z-sections were acquired using Volocity software (Quorum Technologies, Puslinch, ON, Canada). Images were deconvolved (confidence limit of 100% and iteration limit of 30-35) and analyzed with a measurement protocol generated in Volocity (Quorum Technologies), which measured objects in the 3D reconstructed volumes of the optical z-slices. In the measurement protocol described in [11; 108], the PV was pinpointed using the fluorescence intensity of TgGRA7 and TgNTPase; the thresholds of the intensity values were set manually by using the outer TgGRA7 PVM staining as the boundary of the PV. GFP-Rab11A foci were identified by fluorescence intensity (manual thresholds) and the GFP-Rab11A foci located within the boundary of the PV were identified. To compare samples to the control, we used a one-way ANOVA with a Tukey’s honest significant difference post-hoc test (Graphpad Prism).

### Chronic Infections

Female C57BL/6 mice 3-4 weeks old were i.p. infected with 500 or 2500 mutant or wild type Prugniaud*Δ*Ku80 strain tachyzoites harvested from an overnight infected HFF monolayer by mechanical, needle lysis, filtration through a 3 μm nylon filter, washed once with, and resuspended in 1xPBS. Mice were monitored daily for weight and signs of illness. Groups of 4 mice per experiment were used and experiments repeated twice, unless noted otherwise.

Between 3 and 4 weeks post infection, mice were sacrificed through CO2 inhalation, the brains harvested and the cysts enriched and quantified following published methods [109]. In brief, brains were ground up in 1300 μl 1xPBS using mortar and pestle. A 250 μl aliquot of the slurry from two pooled mice brains was subsequently passed five times each through 16G, 18G, 20G and 23G needles. Samples were fixed by adding 150 μl 3% formaldehyde in 1xPBS for 20 min at RT, spun for 5 min at 3000xg and quenched with 150 μl 0.1 M glycine in 1xPBS for 5 min followed by another spin and combined blocking and permeabilization using 150 μl BP-mix (3% BSA in 1xPBS in 0.2% TX-100 in 1xPBS) for 1 hr at RT, or overnight at 4°C. Following a spin, 150 μl of Fluorescein-conjugated *Dolichos bifloris* agglutin lectin (Fluorescein-DBA; 1:3000; Vector Laboratories, FL) in BP-mix was incubated for 1 hr at RT. After three washes with 150 μl BP-mix, 5 μl of the brain pellet was spread and mounted on three different slides, which were all counted by fluorescence microscopy. The total cyst number multiplied by 26 represents the total number of cysts/ single brain. Animal protocols were reviewed and approved by the Boston College IACUC with protocol number 2018-001.

## Rosette assay

Directly A488 conjugated T41E5 *α*-SAG1 antibody [110] kindly provided by Dr. Jean-François Dubremetz was used to stain 4%PFA fixed vacuoles permeabilized with 0.25% TX-100 (both in 1xPBS) 30 hr infected in HFF cells. Vacuoles containing at least 16-parasites were visualized by fluorescence microscopy and enumerated for random or rosette organization of the tachyzoites. Three biological replicates were performed and at least 100 vacuoles per sample were counted.

## 13 Author contributions

M.J.G. designed experiments, generated schematics, wrote the manuscript, acquired funding

D.J.P.F. performed TEM for the BC development cycle

S.S. generated the CaM-cKD line and evaluated the genotype and phenotype

J.D.R. performed host Rab11A-GFP accumulation assays

S.C. generated BC gene knock out strains in the Pru line, performed chronic infection experiments

V.A.P. generated the MyoI-KO parasite line and evaluated phenotype

C.M. generated MyoI-cKD parasite line and performed rosette formation assay on BC mutant strains

I.C. performed TEM on BC mutant strains

K.E. designed experiments, generated BC knock out strains in the RH line, edited the manuscript, acquired funding

All authors proofread the manuscript

## Acknowledgements

We thank Irem Özkan, Karen Zhu, and Nicholas Lo for technical assistance, Bret Judson and the Boston College Imaging Core for infrastructure and support, Drs. David Bzik, Vern Carruthers, Wassim Daher, Chris de Graffenried, Jean-François Dubremetz, and Boris Striepen for sharing reagents and Dr. Tim Gilberger for sharing the TgBTP1 ortholog identity.

## Supplementary Material

**Table S1. Primers used for cloning and PCR validation of transgenic parasites.**

**Supplementary Figure S1.**
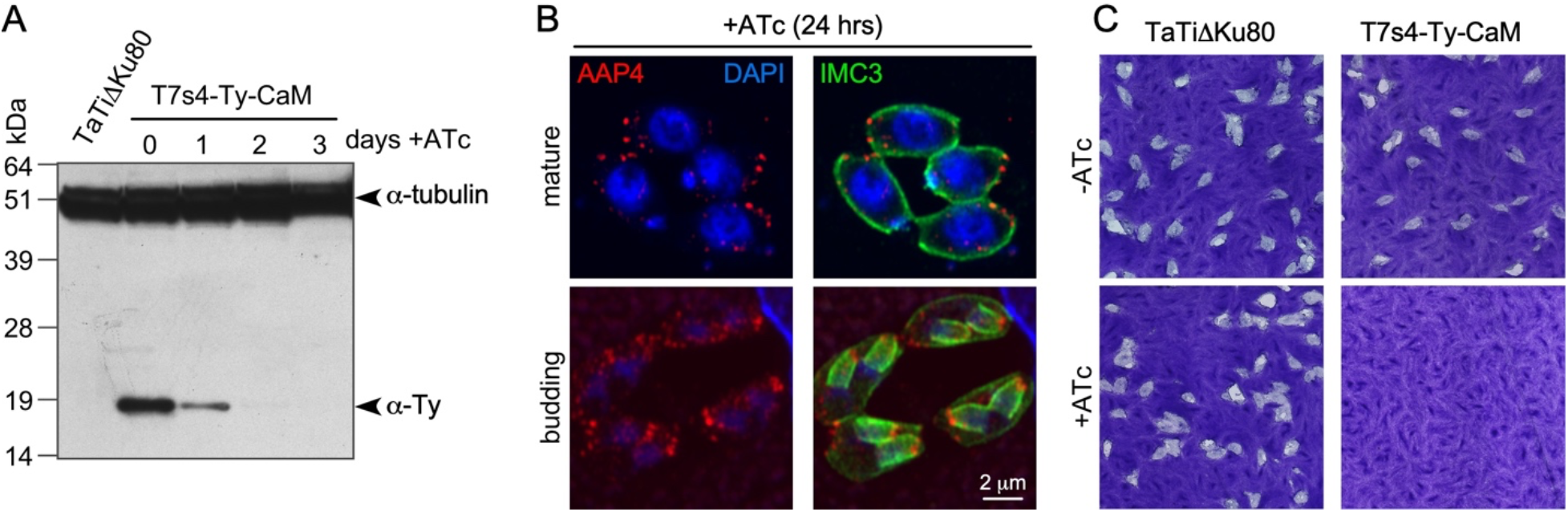
Conditional knock-down of calmodulin does not cause a BC phenotype but is ultimately lethal. A. Western blot analysis displaying the kinetics of CaM depletion upon ATc treatment. Ty highlights CaM, whereas tubulin is used as loading control. B. Depletion of CaM upon ATc addition for 24 hrs does not have an appreciable effect on the morphology of either mother or daughter buds. AAP4 stains the apical annuli and its position marks the apical end or parasites thereby permitting the inference of BC localization at the other end of the cytoskeleton scaffolds marked by IMC3; DAPI marks the DNA. C. Prolonged CaM depletion is lethal as illustrated by plaque assays for 7 days. TaTi*Δ*Ku80 is the parent line.

**Supplementary Figure S2.**
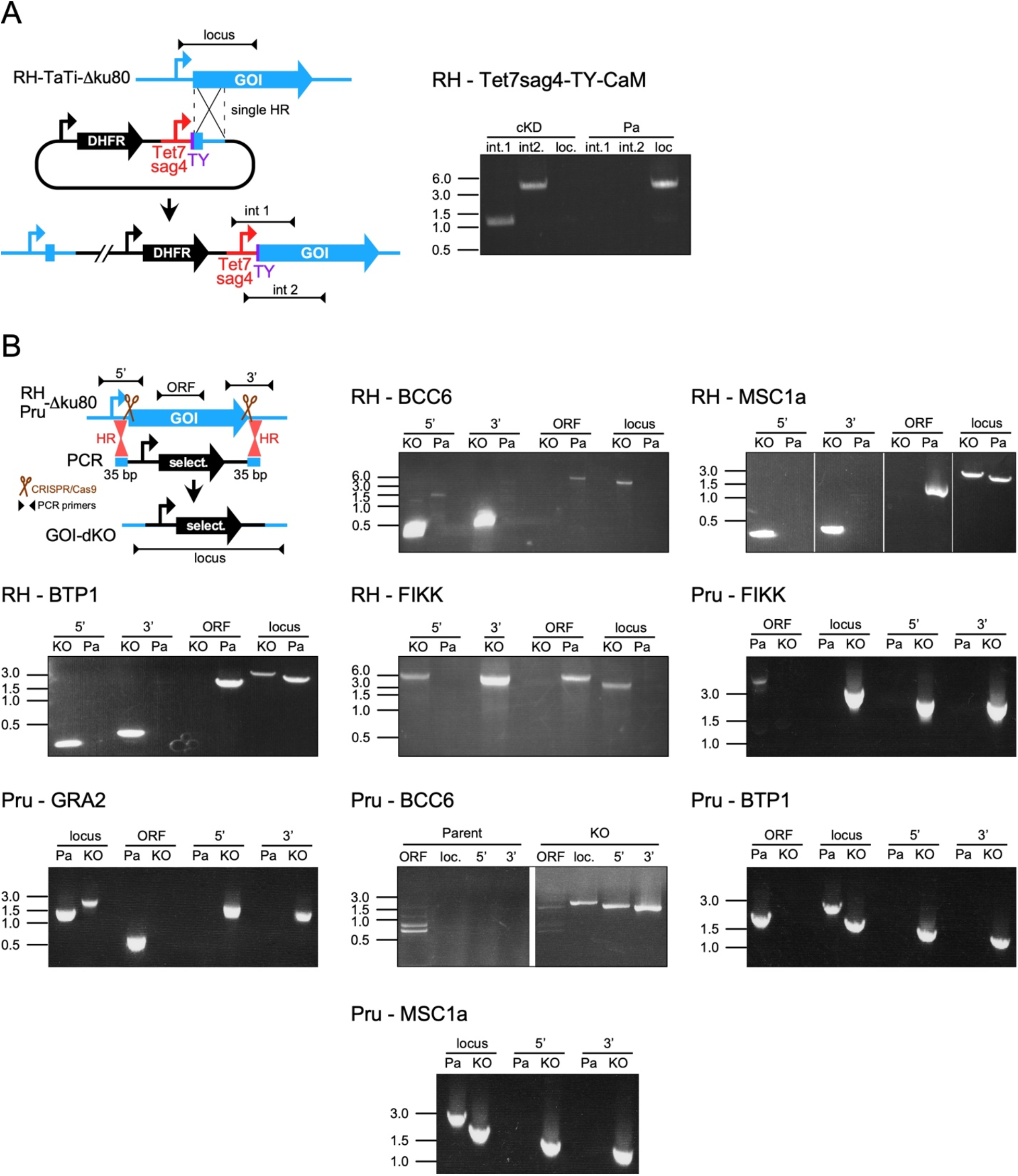
PCR validation of transgenic lines generated in this study. A. Calmodulin conditional knock-down line. cKD, conditional knock-down; locus, the original locus across the targeted site of insertion. int.1 integration primer pair #1; int.2 integration primer pair #2. DNA sizes indicated on the left of all gels. B. knock-out lines, as indicated. select.: drug selectable marker, which was DHFR-TS for RH genotype lines, and HXGPRT for Prugniaud (Pru) genotype lines. KO, knock-out, Pa, parent line; 5’, 5’-flank of the integration site; 3’, 3’-flank of the integration site; ORF, open reading frame of the targeted gene; locus, PCR across the integrated drug selectable marker. DNA sizes indicated on the left of all gels.

